# Visual memories are stored along a compressed timeline

**DOI:** 10.1101/101295

**Authors:** Inder Singh, Aude Oliva, Marc W. Howard

## Abstract

In continuous recognition the recency effect manifests as a decrease in accuracy and a sublinear increase in response time (RT) with the lag of a repeated stimulus. The recency effect could result from the gradual weakening of mnemonic traces. Alternatively, the recency effect could result from a search through a compressed timeline of recent experience. These two hypotheses make very different predictions about the shape of response time distributions. Using highly-memorable pictures to mitigate changes in accuracy enabled a detailed examination of the effect of recency on retrieval dynamics. The recency at which pictures were repeated ranged over two orders of magnitude across three experiments. Analysis of the RT distributions showed that the time at which memories became accessible changed with the recency of the probe, as predicted by a serial search model suggesting that visual memories can be accessed by sequentially scanning along a compressed representation of the past.

---

In recognition memory experiments participants must determine whether a probe stimulus has been previously experienced or not. As the recency of a repeated probe decreases, the accuracy of the judgment decreases and response time (RT) increases (Donkin & Nosofsky, 2012; Hockley, 1982; Monsell, 1978; Murdock & Anderson, 1975; Shepard & Teghtsoonian, 1961). Despite decades of empirical and modeling work, it remains unclear what changes in the state of memory cause the recency effect in recognition memory.

In continuous recognition, an item is presented at each time step; the participant indicates whether it was previously experienced. Because previously-experienced items must be identified from a stream of information, continuous recognition is somewhat similar to the experience of memory in the real world. Consider the task of an individual engaged in continuous recognition (Figure 1a). In order to correctly identify an item as old, one must compare it to the contents of their memory. In continuous recognition, the recency effect manifests as a sublinear increase in RT with increasing lag of the repeated probe (Hintzman, 1969; Hockley, 1982; Okada, 1971); Hockley (1982) found a logarithmic increase in RT with increasing lag. Why does it take longer to retrieve memories from further in the past?

**Figure 1.**
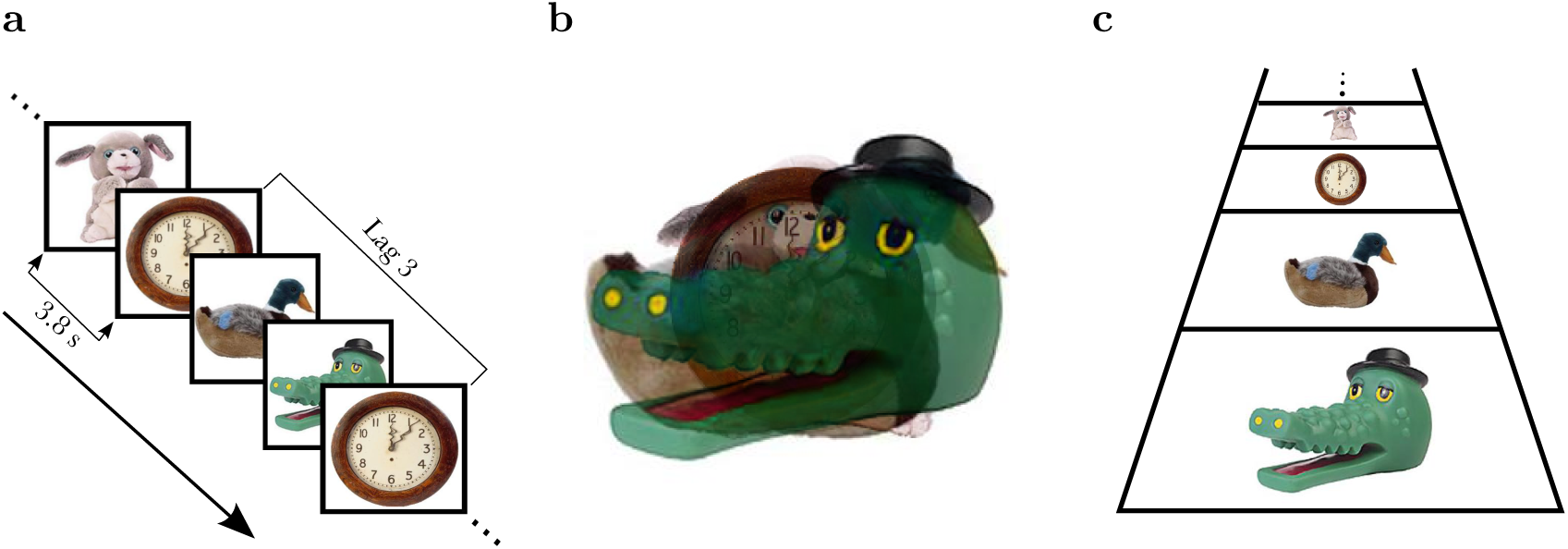
Two forms of memory representations that account for the recency effect. **a.** Schematic of a continuous recognition experiment. Participants experience a long sequence of pictures. Their task is to detect occasional repeated pictures. The variable “lag” measures the difference in position between the two presentations of the repeated picture. **b.** Cartoon representing a composite memory representation. Traces of more recent items (alligator) are more prominent than the traces of less recent items (clock). **c.** Cartoon representing a compressed timeline. This form of representation contains information about the order in which items were experienced. Because the timeline is ordered it can be serially scanned starting at the present (alligator) and going back through the past. The foreshortening of the image is intended to suggest compression.

Many distributed memory models assume that memory is a composite store containing a noisy record of features from all the studied items (e.g., Anderson, 1973; Murdock, 1982; Shiffrin, Ratcliff, Murnane, & Nobel, 1993). A composite memory store can account for the recency effect if the features of items experienced further in the past are stored with less fidelity than items experienced more recently (Figure 1b). In contrast to the “bag of features” of a composite memory, another class of models proposes that features are stored along a timeline of experience (Figure 1c; G. D. A. Brown, Neath, & Chater, 2007; Murdock, 1974; Howard, Shankar, Aue, & Criss, 2015). As an analogy, the memory store behaves like a conveyor belt that recedes into the past (Murdock, 1974). As each item is presented, it is placed at the front of the belt; previously-stored items shift back towards the past.^1^ Both frameworks can accommodate the sub-linear increase in RTs with lag. A composite memory can account for the sub-linear increase if the strength of the match between a probe and the contents of memory decreases appropriately and this strength is coupled with a model of information accumulation (e.g., S. Brown & Heathcote, 2005; Ratcliff, 1978; Usher & McClelland, 2001). Information along a timeline can be sequentially accessed during memory search, which terminates when a match to the probe is found. Critically, if this timeline is compressed into a logarithmic scale (G. D. A. Brown et al., 2007; Chater & Brown, 2008; Shankar & Howard, 2013; Howard et al., 2015), then a logarithmic increase in RT with lag naturally results from this self-terminating search. On a logarithmic scale the difference between lag 1 and lag 2 is larger than the difference between lag 100 and lag 101. Rather, the difference between 1 and 2 is equivalent to the difference between 100 and 200. This property naturally results in a sublinear increase in RT for probes experienced further in the past.

While these two models cannot be distinguished based on RT alone, they make very different predictions regarding the *shape* of the RT distributions (Figure 2). Because all of the traces are stored together in a single composite representation (Figure 2a), the time needed to access a composite memory should be the same regardless of how far in the past the probe was experienced. However, the strength of that match should depend on the probe’s recency. This is analogous to changing the drift rate in a drift diffusion process with increasing lag (Donkin & Nosofsky, 2012; Ratcliff, 1978). A composite memory representation suggests a parallel access model in which the RT distributions rise from zero at the same time but differ systematically in the tail of the distribution as a function of recency. If memories are aligned along a timeline very different qualitative predictions are possible. If the timeline is accessed via a self-terminating serial scan, then the time it takes to access the right memory should depend on the recency of the probe stimulus. Thus this kind of sequential access model results in RT distributions that start at different times (Figure 2b). Although the rate of information accumulation may also depend on lag in a scanning model, a change in the time to initiate the search with lag is a distinctive prediction of a scanning model.

**Figure 2.**
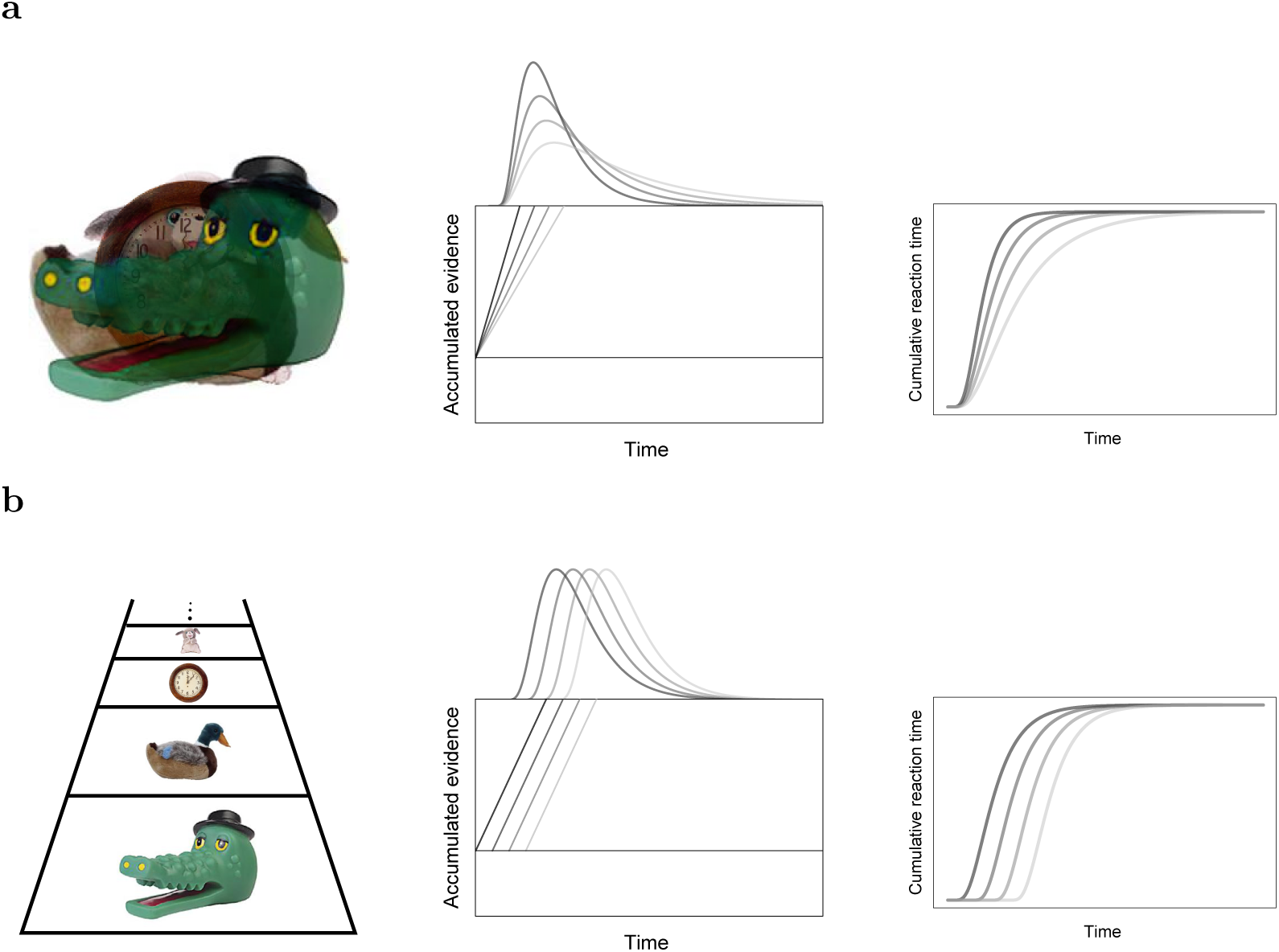
Distinguishing the two forms of memory representation. **a.** A composite memory implies that the items are accessed in parallel. The rate of information accumulation is higher for more recent probes and the time to start accessing the memory of the probe item does not depend on its lag. Thus a parallel access hypothesis results in RT distributions that rise at about the same time but show systematically longer tails as the probe becomes less recent. This trend can be seen readily in cumulative RT plots (right). **b.** A timeline can accommodate a self-terminating scanning hypothesis. Under this hypothesis, more recent probes do not differ in their rate of information accumulation, but rather the time at which information starts to accumulate. Thus a scanning hypothesis results in RT distributions that rise at different times but maintain the same shape.

To the best of our knowledge, a systematic change in the time to initiate the memory search has not been observed in continuous recognition. The main issue in continuous recognition is that as lag increases, accuracy decreases (Hockley, 1982; Shepard & Teghtsoonian, 1961), making it more difficult to measure the effect of recency on retrieval dynamics independently of changes in accuracy. Brady, Konkle, Alvarez, and Oliva (2008) showed participants hundreds of memorable images in a continuous recognition task with lags varying over more than two orders of magnitude, with lags from 1 (no intervening items) to 128. Because the pictures were highly memorable, there was little variability in accuracy, even at very long lags. In addition the RT data are minimally affected by sequential dependencies, which are known to affect RTs in recognition memory (Malmberg & Annis, 2012). Because of the use of highly memorable pictures, wide range of lags tested, and the elimination of sequential dependencies, the Brady et al. (2008) is well-suited to study the effect of recency on RT distributions. This paper analyzes the RT data collected during the Brady et al. (2008) task (referred to as Experiment 1) and replicated the basic finding twice. Experiment 2 was a straightforward replication; In Experiment 3 a visual mask separated the images and studied a wider range of lags.

## Materials and Methods

We analyzed the data collected as a part of the repeat detection task in the visual long term memory experiment conducted by Brady et al. (2008) using mTurk (Experiment 1) and we also did two in lab replications of the same task (Experiments 2 & 3). The main difference between Experiments 2 & 3 is that we used a mask to separate the images and added repetitions at longer longs in Experiment 3. Categorically distinct images were obtained from a commercially available database (Hemera Photo-Objects,Vol. I and II) and through internet searches using Google Image Search. Examples of the images are used in Figure 1. The images (subtending approximately 7.5° by 7.5° of visual angle) were presented one at a time at the center of the screen. Participants were required to respond to repeated images but not required to respond “no” to new items. The sequence of presentations did not include successive repetitions. As a result, sequential responses were only made when a repetition was followed by a false alarm or vice versa. Because the false alarm rate was very low, this was quite rare.

Lag was defined as the difference between the position of a repetition and the previous presentation of that stimulus; immediate repetition thus corresponds to a lag of 1. In Experiment 1, we analyzed memory for repeated pictures that were presented within the same block at lags from 1 to 128.^2^ The way the experiment was constructed, repetitions at longer lags were somewhat less likely than repetitions at shorter lags. The average number of observations per participant included in our main analyses ranged from 27 at lag 1 to 9 at lag 128 for Experiment 1. In Experiment 2 the average number of observations per participant varied from 30 at lag 1 to 16 at lag 128 and in Experiment 3 the average number of observations per participant varied from 30 at lag 1 to 12 at lag 512.

There is strong evidence suggesting that the time to access immediate repetitions is very different from the time to access repetitions after intervening stimuli, (e.g., McElree & Dosher, 1989). In order to ensure that the findings were not driven by immediate repetitions, we excluded lag 1 from the analyses. Noting the sublinear effect of lag on RT, we perform statistical analyses on the base 2 logarithm of lag. That is, for the purpose of statistical analyses lags 2, 4, 8, 16…are given values 1, 2, 3, 4…. This means that when a regression coefficient is reported for lag, it is interpretable as the change associated with doubling of lag. For simplicity of exposition, in some cases we will not explicitly state “base 2 log of lag” but simply refer to “lag” in describing statistical analyses.

Some of the images were repeated multiple times during the course of the experiment. Only the first repetitions are included in the first round of analyses. Second repetitions are analyzed later in the subsection entitled “Analysis of second repetitions”. The specific differences between the three experiments are listed below.

### Experiment 1

The accuracy data from Experiment 1 were originally reported in Brady et al. (2008), which provides a detailed description of the methods. During this task a total of 2896 images (2,500 new and 396 repeated images) were shown to 14 participants across 10 study blocks of approximately 20 minutes each. The images were presented for 3 s each, followed by an 800 ms fixation cross.

### Experiment 2 & 3

Experiments 2 and 3 were implemented in PyEPL (Geller, Schlefer, Sederberg, Jacobs, & Kahana, 2007). Participants from Boston University were recruited for one session each and were paid $15 per hour for their time. In Experiment 2, a total of 900 images (650 new and 250 repeated images) were shown to 35 participants across 2 study blocks of approximately 20 minutes each. In Experiment 3, a total of 1360 images (1084 new and 276 repeated images) were shown to 39 participants across 2 study blocks of approximately 35 minutes each. The images were presented for 2.6 s each, followed by a 400 ms of cross for Experiment 2. In Experiment 3 the images were separated by phase scrambled images as masked separators instead of the cross.

### Model fitting

In addition to standard distributional measures, we also characterized RT distributions using the shifted Wald distribution. The shifted Wald distribution gives the finishing times for a drift diffusion process (Wald, 1947) with one absorbing boundary. This is appropriate for this dataset because the participants only provided “yes” responses. The density function of the shifted Wald distribution is described by three parameters, *μ*, λ, and *σ* and is given by:

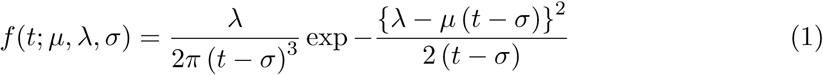

The parameter *μ* describes the rate of information accumulation, i.e., the drift rate. The parameter λ describes the distance of the boundary from the starting point of the diffusion process and *σ* describes the non-decision time before information begins to accumulate.

To determine if there was a significant effect of the time at which information accumulation begins, we considered a model where *σ* and *μ* were allowed to vary freely as a function of lag.^3^ Since we are fitting three parameters per participant for each lag, we only included the subjects who had at least four correct responses per lag. All participants in Experiments 1 and 2 met this threshold. In Experiment 3, this cut-off was met by 32 out of 39 participants.

Log likelihood of each response was computed using the analytic expression in Eq. 1 for each participant assuming responses are independent. The optimization function tried to minimize the negative log likelihood of the data given the parameters varying for each model using the Nelder-Mead algorithm. To avoid local minima, each parameter optimization was run from multiple starting points and the parameters from one iteration were passed back to the optimizer until the parameter values stopped changing.

## Results

### There was a recency effect on accuracy, but hit rate remained high at all lags

In all three experiments, accuracy decreased with increasing lag, but remained quite high (Figure 3a). In Experiment 1, the overall hit rate was .95, compared to a false alarm rate (incorrect detection of new images) of .01, corresponding to a *d*′ of about 4. Hit rate decreased with lag; at lag 128, the hit rate was .89 corresponding to a *d*′ of about 3.5. In Experiment 2, the overall hit rate was .89 and the false alarm rate was .03, corresponding to a *d*′ of about 3. The hit rate varied from 0.96 (*d*′ = 3.57) for immediate repetitions to .69 (*d*′ = 2.3) for a lag of 128. In Experiment 3 the overall hit rate was .81 and the false alarm rate was .03, corresponding to a *d*′ of about 2.8. The hit rate for a lag 1 was 0.92 (*d*′ = 3.28) and dropped to 0.57 (*d*′ = 1.9) for lag 512. Overall, the base 2 logarithm of lag was a significant predictor of the hit rate, across the three experiments (see slope of Hit rates in Table 1). This drop in hit rate seemed to accelerate at longer lags.

**Figure 3.**
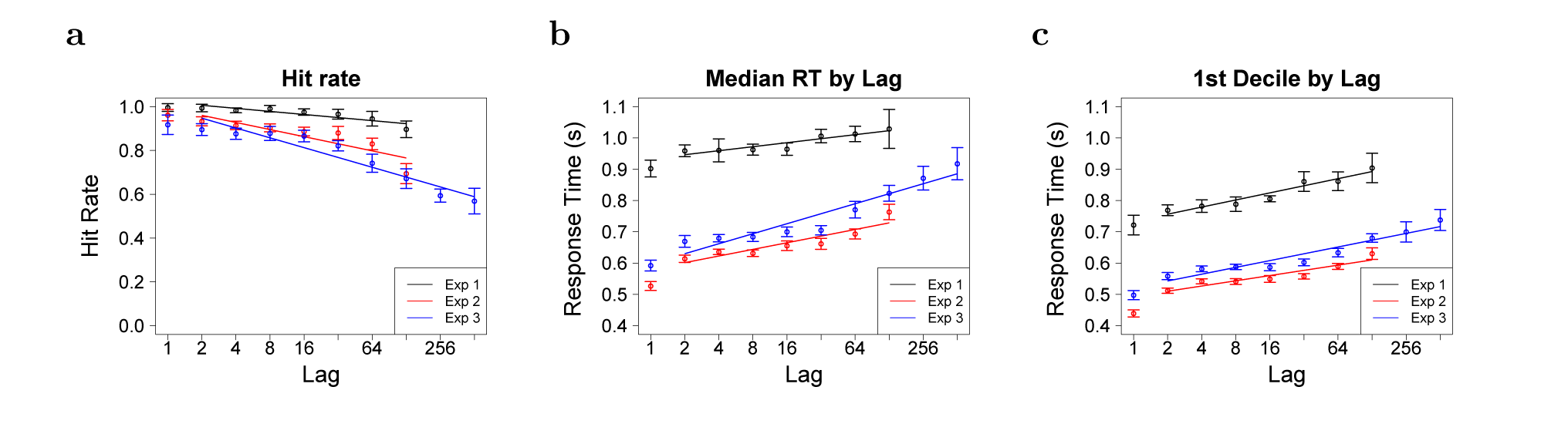
**a.** Hit Rate as a function of lag on log_2_ paper for Experiments 1, 2, and 3. The hit rate goes down with lag. There appears to be a more pronounced drop in the hit rate at higher lags. **b.** Median response time as a function of lag on log_2_ paper. Median response time increased approximately linearly with the logarithm of lag. c. The 1^st^ decile as a function of lag on log_2_ paper. Error bars in all figures represent the 95% confidence interval of the mean across participants normalized using the method described in Morey (2008).

**Table 1:** The slopes and intercepts of the empirical and model fits across the three experiments. Statistical tests were done on the base 2 logarithm of lag; regression coefficients with respect to lag are understandable as the rate per doubling of lag. Effects significant at *p* < 0.001 are indicated by **.

|  |  | Experiment 1 | Experiment 2 | Experiment 3 |
| --- | --- | --- | --- | --- |
| Hit Rate | Intercept | $1.02 \pm 0.01$<br>$t(83) = 82.12^{**}$ | $0.99 \pm 0.02$<br>$t(209) = 48.64^{**}$ | $0.99 \pm 0.02$<br>$t(255) = 43.89^{**}$ |
| | Slope | $-0.014 \pm 0.002$<br>$t(83) = -6.46^{**}$ | $-0.032 \pm 0.003$<br>$t(209) = -11.46^{**}$ | $-0.045 \pm 0.002$<br>$t(255) = -18.62^{**}$ |
| Median | Intercept | $0.93 \pm 0.04$<br>$t(83) = 22.22^{**}$ | $0.58 \pm 0.01$<br>$t(209) = 43.87^{**}$ | $0.598 \pm 0.02$<br>$t(255) = 30.61^{**}$ |
| | Slope | $0.013 \pm 0.003$<br>$t(83) = 4.54^{**}$ | $0.021 \pm 0.002$<br>$t(209) = 13.27^{**}$ | $0.032 \pm 0.002$<br>$t(255) = 17.52^{**}$ |
| 1st decile | Intercept | $0.73 \pm 0.03$<br>$t(83) = 22.05^{**}$ | $0.493 \pm 0.008$<br>$t(209) = 59.28^{**}$ | $0.52 \pm 0.01$<br>$t(255) = 37.43^{**}$ |
| | Slope | $0.023 \pm 0.002$<br>$t(83) = 9.29^{**}$ | $0.017 \pm 0.001$<br>$t(209) = 15.21^{**}$ | $0.022 \pm 0.001$<br>$t(255) = 17.65^{**}$ |
| Shift | Intercept | $0.37 \pm 0.06$<br>$t(83) = 6.72^{**}$ | $0.394 \pm 0.009$<br>$t(209) = 42.21^{**}$ | $0.41 \pm 0.01$<br>$t(255) = 33^{**}$ |
| | Slope | $0.022 \pm 0.003$<br>$t(83) = 6.66^{**}$ | $0.015 \pm 0.001$<br>$t(209) = 13.28^{**}$ | $0.019 \pm 0.002$<br>$t(255) = 12.67^{**}$ |
| Drift | Intercept | $0.57 \pm 0.07$<br>$t(83) = 7.91^{**}$ | $0.23 \pm 0.02$<br>$t(209) = 14.74^{**}$ | $0.25 \pm 0.02$<br>$t(255) = 11.41^{**}$ |
| | Slope | $-0.003 \pm 0.005$<br>$t(83) = -0.72, ns$ | $0.011 \pm 0.002$<br>$t(209) = 4.46^{**}$ | $0.014 \pm 0.002$<br>$t(255) = 6.23^{**}$ |
| Shift-<br>till lag 32 | Intercept | $0.36 \pm 0.06$<br>$t(55) = 6.57^{**}$ | $0.401 \pm 0.009$<br>$t(139) = 44.2^{**}$ | $0.43 \pm 0.01$<br>$t(127) = 40.35^{**}$ |
| | Slope | $0.024 \pm 0.005$<br>$t(55) = 5.15^{**}$ | $0.012 \pm 0.002$<br>$t(139) = 7.79^{**}$ | $0.011 \pm 0.002$<br>$t(127) = 5.14^{**}$ |
| Drift-<br>till lag 32 | Intercept | $0.59 \pm 0.07$<br>$t(55) = 8.23^{**}$ | $0.24 \pm 0.02$<br>$t(139) = 14.89^{**}$ | $0.28 \pm 0.02$<br>$t(127) = 12.16^{**}$ |
| | Slope | $-0.011 \pm 0.006$<br>$t(55) = -1.94$ | $0.004 \pm 0.003$<br>$t(139) = 1.37$ | $0 \pm 0.004$<br>$t(127) = 0.06$ |

### Non-parametric measures of RT distributions showed that more recent memories were accessed more quickly

The median RT increased as a function of (base 2 logarithm of) lag in all three experiments (Figure 3b). Allowing for independent intercepts for each participant using linear mixed effects model, we found that lag was a significant predictor of the median response time across Experiment 1, 2 & 3 (Table 1).

A serial self-terminating model predicts RT distributions that change systematically with lag in terms of their starting positions (Figure 2b). This implies that the difference due to lag should be observable at the earliest parts of the distribution. Examination of the cumulative RT distributions (Figure 4) appeared to show a systematic change in the starting position of the distributions as a function of lag across all three experiments. In order to quantify this visual impression, we analyzed the 1^st^ decile of the response time distribution. Across the three experiments, the slope of the 1^st^ decile was significant. Notably, the slope of the 1^st^ decile was comparable over the three experiments, with overlapping confidence intervals in Experiments 1 & 3. The value of the slope indicates that each doubling of lag resulted in a shift of about 20 ms in the distribution.

**Figure 4.**
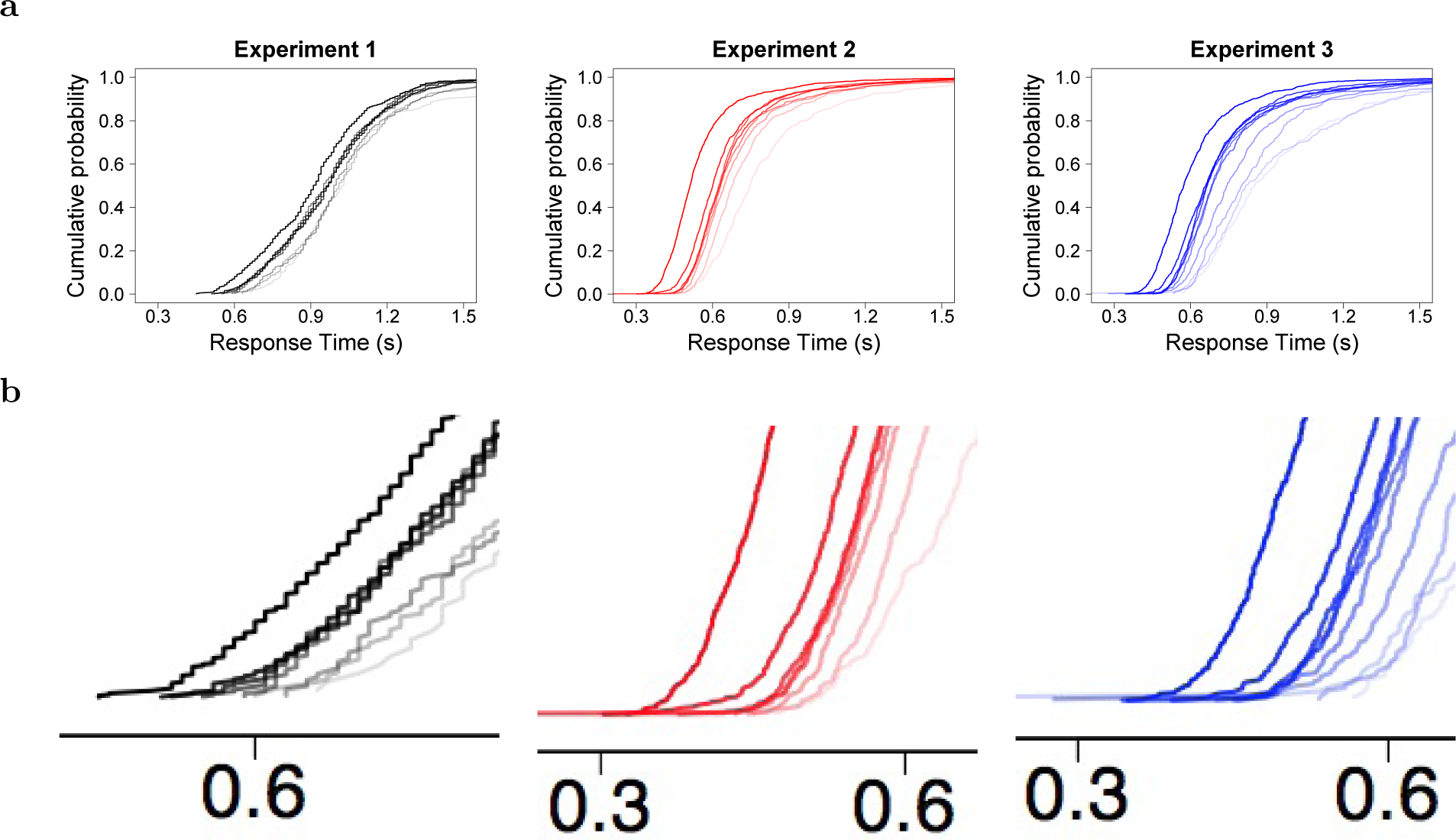
Time to access memory changed systematically with lag in all three experiments. **a.** Unsmoothed across-participant cumulative response distributions for each lag. Shorter lags are darker shades. Note that the cumulative distributions shift with decreasing recency. The three panels are Experiments 1-3 (left to right). Compare to Figure 2 (right). **b.** Inset to make the shift more readily apparent.

### Model-based analyses of RT distributions showed that more recent memories were accessed more quickly

The model-based analyses of RT distributions confirmed the impression from non-parametric statistics that the RT distributions shifted with increasing lag. Figure 5 summarizes the results for the shift and drift parameters of the shifted Wald distribution; statistics are shown in Table 1. To summarize the results, the shift parameter showed a consistent effect of lag across all three experiments. The intercept for the shift parameter, about 400 ms, was similar for the three experiments despite large changes in median RT. All three experiments showed reliable slopes demonstrating that the shift parameter *σ* changed systematically as a function of lag. The regression coefficients were similar for the three experiments (the confidence intervals for Experiments 1 and 3 overlapped); each doubling of lag shifted the time to access memory by about 15-20 ms.

**Figure 5.**
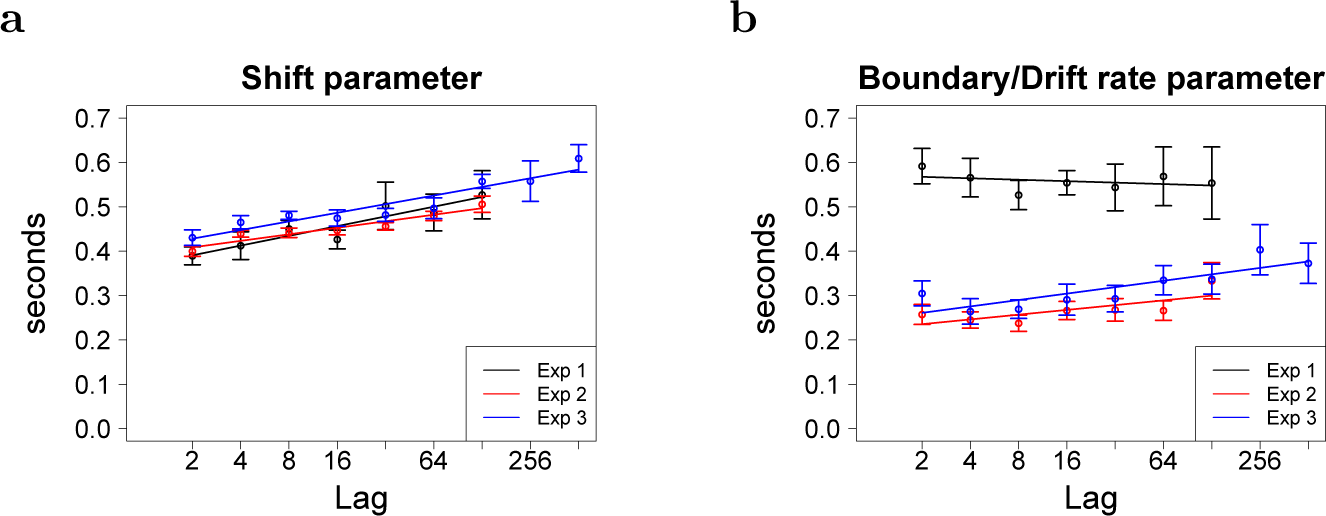
The time to access memory, estimated by the shift parameter of the Wald distribution, changed systematically with lag and was consistent across the three experiments. **a.** The shift parameter of the Wald distribution as a function of lag on log2 paper for the three experiments. **b.** The drift parameter of the Wald distribution as a function of lag across the three experiments.

To assess the effect of lag on μ in similar units, we took the λ divided by *μ*^4^ as a dependent measure. In contrast the drift parameter μ did not show a consistent trend across experiments (see Figure 5).

Qualitatively, it seems that the drift parameter did not change reliably at relatively short lags. In order to assess this, noting that lag 32 was the point at which hit rate began to fall off more abruptly (Fig. 3a), we recomputed statistics for shift and drift for lags 2-32. The results are in Table 1. Over this range of lags, the shift parameter still showed a reliable linear trend, whereas the drift parameter did not. Moreover, the confidence intervals for the regression coefficients did not overlap in any of the three experiments, demonstrating that for lags from 2 to 32 there was a differential effect of recency on the shift parameter *vs* the drift parameter. Over this range of lags, corresponding to delays of up to about a minute and a half, the effect of recency was solely carried by the shift parameter.

We find converging evidence in favor of a systematic change in the non-decision time with lag from the model based analyses. The shift parameter has a significant slope across the three experiments. Thus the evidence from non-parametric analyses and model-based analyses strongly support the hypothesis that the time necessary to access visual memories changes with recency.

### Analysis of second repetitions argued against a non-scanning account of the results

Analysis of the first repetitions showed evidence that recency affects RT distributions *via* a shift in the distribution, consistent with a change in the time to access memory as predicted by a self-terminating search model. While a shift in the RT distribution is consistent with a serial self-terminating search during the memory comparison phase, this is not the only possible explanation. Presumably, the probe must be encoded before it can be compared to memory. The shift in the RT distributions is also consistent with the hypothesis that encoding of recently-experienced probes is facilitated but that there is no effect of recency on the memory-comparison stage *per se* (Sternberg, 1969).

If the recency of a repeated item allows it to be processed faster as a probe, then repeating the item again should have an additional effect on RT. Thus far we have examined RTs to the first repetition (second presentation) of the probe stimulus. In order to evaluate the hypothesis that the recency effect was attributable to processing fluency, we examined RTs to the second repetition (third overall presentation). If changes in RTs were being driven by greater facility of processing of recently-presented probes, then the lags of the two presentations ought to both affect RT. In contrast, if the changes in RT are driven by a self-terminating scanning model during the memory comparison stage, then only the most recent lag should affect RT. Scanning further predicts that the effect of the most recent lag should be the same on both the first and second presentations and that in both cases the effect of lag should be to cause a shift of the RT distribution. All three of the predictions of a self-terminating scanning model held across all three experiments.

Accuracy was very high for probes that were repeated a second time. In Experiment 1, 49 images that were repeated a second time and the overall hit rate for the second repetitions was 0.99 ± 0.006. In Experiments 2 and 3, 54 images were presented a third time and the overall hit rates were 0.98 ± 0.01 and 0.96 ± 0.01 respectively.

#### Only the most recent lag affected RT on second repetitions

There are two lags associated with the third presentation of a probe. Let us refer to the lag between the first and second presentation as lag_1_ and the lag between the second and third presentation as lag_2_, so that at the third presentation lag_2_ is the most recent lag. Allowing each participant to have an independent intercept (to account for between participant variability) in a linear mixed effects model using the base 2 logarithm of the two lags as regressors, there were reliable effects of lag_2_ but no effect of lag_1_ in all three experiments. In Experiment 1 lag_2_ showed a reliable effect, .016 ± .005, *t*(600) = 3.39, *p* < .01; but no effect of lag_1_, –.003 ± .005, *t*(600) = –.60. In Experiment 2, lag_2_ showed a reliable effect, .012 ± .003, *t*(1812) = 3.84, *p* < .01, but lag_1_ did not, –.001 ± .003, *t*(1812) = .02. In Experiment 3, the same pattern held, lag_2_: .015 *±* .004, *t*(1623) = 3.44, *p* < .01; lag_1_, –.007 ± .005, *t*(1623) = 1.5.

#### The effect of lag on RT was the same for first and second repetitions

In the analysis shown above, RT to images repeated a third time depended on the most recent lag (second repetition) and did not depend on the lag to the first repetition. A scanning model predicts further that the effect of lag_2_ on the second repetition should be the same as the effect of lag on the initial repetition. Figure 6a-c summarize these comparisons. Across all three experiments, plots of RT as a function of lag (lag_2_ for repeated stimuli) showed parallel curves for first and second repetitions on log2 paper. This visual impression was confirmed by a multiple regression with lag, repetition and an interaction term of lag and repetition. In all the three experiments, there was a significant effect of lag and repetition but the interaction term was not significant (statistics are in Table 2).

**Figure 6.**
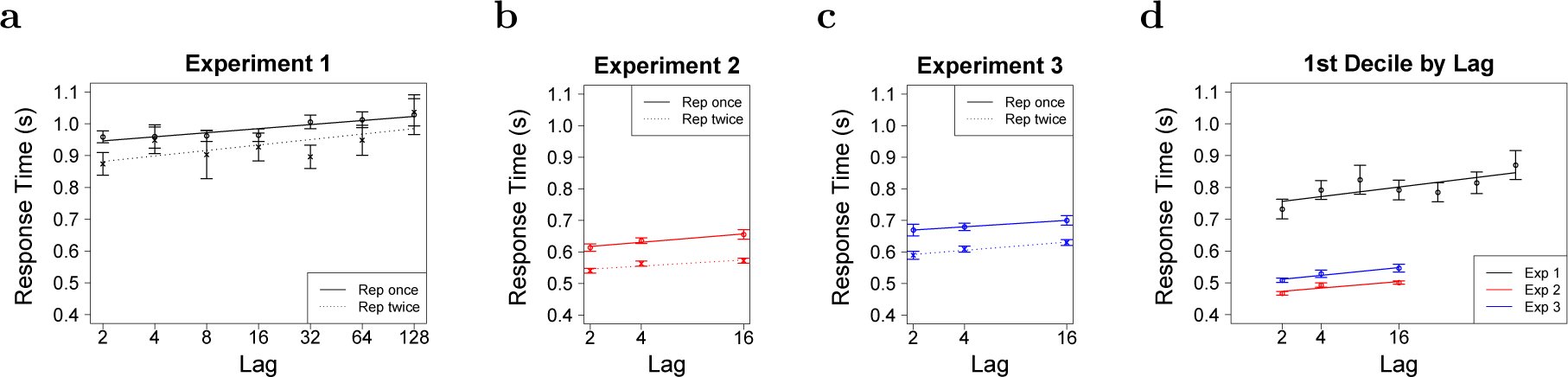
The effect of lag on RT was the same for first and second repetitions. **a-c.** Median response time as a function of the most recent lag for first (solid) and second (repetitions) for the three experiments. To the extent the lines are parallel, it means that the effect of recency on RT was the same on the first and second presentations. Analyses reported in the text demonstrated that only the most recent lag affected the RT of old probes repeated twice. **d.** The 1^st^ decile of the second repetitions as a function of lag on log_2_ paper. The results are comparable to those for first repetitions (Fig. 3c). Distributions for second repetitions are shown in Fig. 7.

**Table 2:**
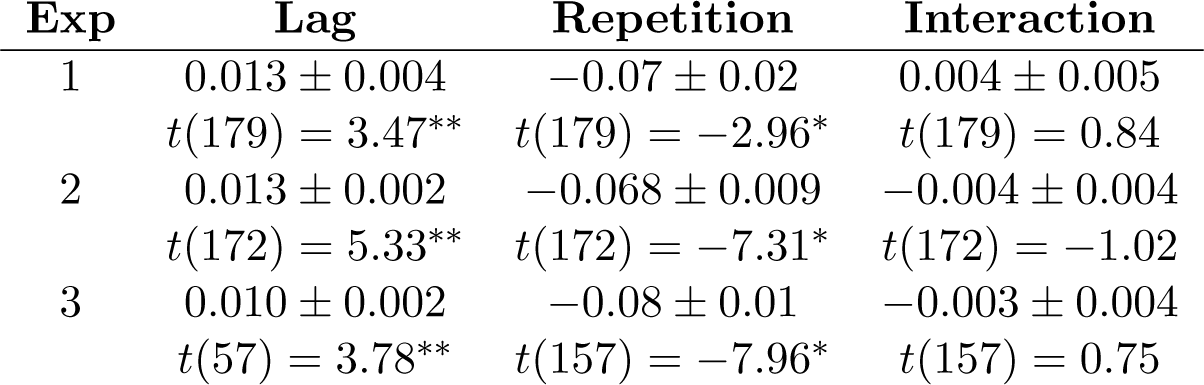
The RTs to second repetitions are faster than to first repetitions but both repetitions show the same effect of lag. Results of a multiple regression analysis with (base 2 logarithm of) lag, repetition and their interaction. See also Figure 6a-c. Effects significant at *p* < 0:001 are indicated by ** and at *p* < 0.01 are indicated by *.

| Exp | Lag | Repetition | Interaction |
| --- | --- | --- | --- |
| 1 | $0.013 \pm 0.004$ | $-0.07 \pm 0.02$ | $0.004 \pm 0.005$ |
| | $t(179) = 3.47^{**}$ | $t(179) = -2.96^*$ | $t(179) = 0.84$ |
| 2 | $0.013 \pm 0.002$ | $-0.068 \pm 0.009$ | $-0.004 \pm 0.004$ |
| | $t(172) = 5.33^{**}$ | $t(172) = -7.31^*$ | $t(172) = -1.02$ |
| 3 | $0.010 \pm 0.002$ | $-0.08 \pm 0.01$ | $-0.003 \pm 0.004$ |
| | $t(57) = 3.78^{**}$ | $t(157) = -7.96^*$ | $t(157) = 0.75$ |

#### RT distributions shifted as a function of recency on second repetitions

A scanning account predicts that the effect of lag on second repetitions should not only be of a similar magnitude as the effect of lag on first repetitions, but that in both cases the effect should be associated with a shift in the distributions. Figure 7 shows the cumulative RT distributions for second repetitions for different lags. As in the initial repetitions (Fig. 4), the RT distributions appeared to shift with lag. This visual impression was conformed by non-parametric analysis.^5^ Figure 6d shows the 1^st^ decile of the RT distributions as a function of lag for all three experiments. A regression of the 1^st^ decile of the RT for the second repetitions onto lag was significant across the three experiments (Exp. 1: 0.015 ± 0.003, *t*(83) = 4.5; *p* < 0.001; Exp. 2: 0.010 ± 0.001, *t*(69) = 6.9, *p* < 0.001; Exp. 3: 0.012 ± 0.002, *t*(63) = 5.08; *p* < 0.001).

**Figure 7.**
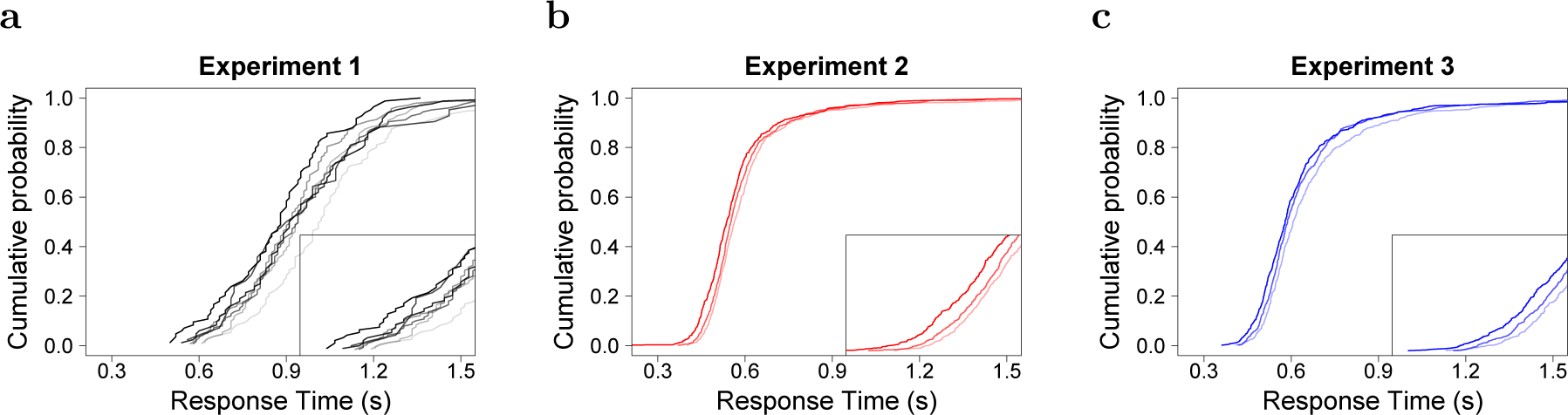
For probes repeated a second time, the most recent lag affected the time to access memory. Unsmoothed across-participant cumulative response distributions for second repetitions are shown for the most recent lag. Shorter lags are darker shades. The inset zooms in on the time at which the distributions start rising from 0. **a.** Experiment 1. **b.** Experiment 2. **c.** Experiment 3.

To summarize, while there was an effect of the lag to the most recent presentation of the probe stimulus, there was no additional effect of its prior presentation. The response time distributions for the second repetitions show a systematic change in the time at which cumulative distributions rise from zero and the effect of multiple repetitions manifests itself in the intercept term of the response time vs. lag line. These observations fail to provide any evidence for the hypothesis that the effect of recency on RT was due to a facilitation of probe encoding. In contrast, these findings are exactly as one would have predicted if the effect of recency on RT was caused by serial self-terminating scanning of a compressed representation of the past.

## Discussion

It has long been known that response time in a continuous recognition experiment increases with the lag to the probe. If memory search requires scanning of a timeline to find the appropriate memory, then the increase in RT should be associated with a shift in the distribution. Three experiments showed that lag consistently shifted RT distributions. This effect was observable both using non-parametric analyses of RT distributions (Fig. 4) as well as a model-based analysis (Fig. 5). Consistent with the scanning account, RTs to second repetitions depended only the most recent lag, as if the search terminated when the first match is found. Although RTs were faster to second repetitions overall, the effect of the most recent lag on second repetition RTs was the same as the effect of lag on first presentation RTs (Fig. 6a-c). This effect on second repetitions was manifest as a shift of the RT distributions (Fig. 7).

Consistent with earlier findings (e.g., Hockley, 1982) the results of these experiments showed a sublinear shift with lag. The results of this paper are roughly consistent with a logarithmic increase in RT as a function of lag; each doubling of lag resulted in a shift of approximately 15-20 ms in the RT distribution. We observed a continuous shift in RT distributions from lags covering a few seconds up to lags of more than 20 minutes. The results of this study are consistent with scanning along a logarithmically-compressed timeline.

There is extensive evidence for self-terminating serial search models in short-term memory tasks (Hacker, 1980; Hockley, 1984; McElree & Dosher, 1993; Sternberg, 2016). There is also evidence consistent with scanning along a timeline in JOR tasks over longer time scales (Hintzman, 2010). However, most previous studies of item recognition have found evidence for parallel access to memory, not sequential scanning, in study-test recognition (Nosofsky, Little, Donkin, & Fific, 2011; Ratcliff & Murdock, 1976; McElree & Dosher, 1989; Hockley, 1984; Nosofsky, Cox, Cao, & Shiffrin, 2014). Several potentially important methodological differences may account for the discrepancy between those studies and the results in this paper. This experiment used continuous recognition rather than the study-test procedure (Nosofsky et al., 2011; Ratcliff & Murdock, 1976; McElree & Dosher, 1989; Hockley, 1984; Nosofsky et al., 2014) and highly memorable trial-unique visual stimuli. In addition, this study only required positive responses to repeated stimuli and more recent lags were tested more frequently than more remote lags. The question of which combination of these methodological differences accounts for the evidence for serial scanning is an extremely important one that merits further investigation. It is worth noting that there is no reason in principle that a compressed timeline could not be accessed in parallel (Howard et al., 2015), whereas it is not clear how (or why) a composite representation could be scanned in a recognition memory task. Notably, if a logarithmically-compressed timeline is accessed in parallel, the recency effect for strength of match would fall off like a power law (Howard et al., 2015), much like the change Donkin and Nosofsky (2012) observed experimentally in drift rate.

On its face, serial scanning of a compressed timeline requires a more elaborate memory representation than a simple composite memory. However, it also suggests a deep analogy between search through memory and directed attention along perceptual dimensions. Neural receptive fields in vision form a compressed representation of retinal space, with broader receptive fields at locations further from the fovea (Hubel & Wiesel, 1974; Schwartz, 1977). By analogy, “time cells” in the hippocampus (MacDonald, Lepage, Eden, & Eichenbaum, 2011), prefrontal cortex (Tiganj, Kim, Jung, & W., in press) and striatum (Mello, Soares, & Paton, 2015) can be understood as constructing a compressed timeline. We can deploy attention strategically to sensory dimensions resulting in preferential access to information available along those dimensions (e.g., Teder-Sälejärvi, Münte, Sperlich, & Hillyard, 1999; Shomstein & Yantis, 2004). To the extent both perception and memory make use of the same form of compressed neural representation, scanning in memory can be understood as exploiting the same kind of computational mechanisms used to direct visual attention.

## Footnotes

1 In this study time *per se* and number of intervening items are confounded so we will not attempt to disentangle them.

2 We excluded lag 256 in Experiment 1, which was repeated fewer than 5 times per participant.

3 Note that it is not sensible to allow boundary separation to vary as a function of recency—this would require that the participant know the recency of the probe before information begins to accumulate.

4 The drift rate *μ* is in units of evidence per unit time. The boundary separation λ is in units of evidence. λ/*μ* thus has units of time, making it directly comparable to σ.

5 Because the number of responses per lag to second repetitions was relatively small, we did not attempt model-based analyses of second repetitions.

